# Transcription-Coupled Repair of DNA Interstrand Crosslinks by UVSSA

**DOI:** 10.1101/2023.05.10.538304

**Authors:** Rowyn C Liebau, Crystal Waters, Arooba Ahmed, Rajesh K Soni, Jean Gautier

## Abstract

DNA interstrand crosslinks (ICLs) are covalent bonds between bases on opposing strands of the DNA helix which prevent DNA melting and subsequent DNA replication or RNA transcription. Here, we show that Ultraviolet Stimulated Scaffold Protein A (UVSSA) participates in transcription-coupled repair of ICLs in human cells. Inactivation of UVSSA sensitizes human cells to ICL-inducing drugs, and delays ICL repair. UVSSA is required for transcription-coupled repair of a single ICL in a fluorescence-based reporter assay. UVSSA localizes to chromatin following ICL damage, and interacts with transcribing Pol II, CSA, CSB, and TFIIH. Specifically, UVSSA interaction with TFIIH is required for ICL repair. Finally, UVSSA expression positively correlates with ICL chemotherapy resistance in human cancer cell lines. Our data strongly suggest that transcription-coupled ICL repair (TC-ICR) is a *bona fide* ICL repair mechanism that contributes to crosslinker drug resistance independently of replication-coupled ICL repair.

## INTRODUCTION

DNA interstrand crosslinks (ICLs) are a particularly genotoxic type of DNA adduct that occurs when a covalent bond forms between bases on opposing strands of the DNA helix. ICLs prevent melting of double stranded DNA, blocking essential DNA transactions including RNA transcription and DNA replication. As a result, these lesions are highly cytotoxic^1,2^. ICLs are caused by endogenous metabolites as well as exposure to chemotherapeutic drugs. Unrepaired ICLs disrupt replication, halt the cell cycle, and trigger apoptosis^3^. To preserve genome integrity, organisms have evolved multiple repair mechanisms to remove ICLs. These pathways can be divided into three categories based on the method of lesion detection: replication-coupled, transcription-coupled, and direct detection.

Replication-coupled repair via the Fanconi Anemia (FA) pathway is the best characterized mechanism for ICL removal. The FA pathway was identified in patients suffering from Fanconi Anemia, an inherited disorder characterized by bone marrow failure, high incidence of early onset cancer, and extreme sensitivity to crosslinking agents. Study of FA patient derived cells has revealed a large group of mutated genes and elucidated the proteins involved in the FA pathway^1,4,5^. FA is activated during DNA replication, when a replication fork stalls at an ICL. The stalled fork recruits FA proteins and the FA core complex, which activate downstream repair mechanisms to promote removal of the lesion and resumption of replication. Many mechanistic insights into these processes were defined using cell free extracts derived from *Xenopus laevis* eggs^6–8^. Cells harboring mutations in the FA pathway are not able to resolve stalled replication forks at ICLs and accumulate DNA double strand breaks (DSBs), mutations, and rearrangements^1,4,5^. Replication-coupled repair by FA is thought to be the major repair mechanism to remove ICLs. However, non-replicative or rarely replicating cells must still repair ICLs to preserve genome integrity. Replication independent repair (RIR) of ICLs, via direct detection or transcription-coupled pathways, provides critical ICL repair mechanisms that function independently of FA^2,9,10^.

Direct detection based ICL repair is initiated by proteins that detect the helix distortion caused by ICLs. Several pathways for direct detection mediated repair have been identified in model systems that undergo RIR. For example, replication-incompetent *Xenopus laevis* cell-free extracts support ICL repair. In this system, RIR requires the DNA polymerase Pol κ, implicating translesion synthesis (TLS) in ICL removal^11^. Upstream ICL sensing in *Xenopus* extract involves the Mismatch Repair (MMR) protein MutSα, a DNA damage sensor that detects helix distortion caused by DNA lesions. MutSα initiates repair of ICLs via MMR, triggering repair independently of DNA transactions^12^. In G1 arrested mammalian cells, the damage sensor XPC is required for ICL repair, revealing another direct detection based ICL repair mechanism^13^ which likely functions via the nucleotide excision repair (NER) pathway^14^.

Finally, transcription-coupled ICL repair (TC-ICR) has been described in several experimental settings. In hamster cells, repair of ICLs is biased towards transcribed regions of the genome^15^. Loss of transcription-coupled repair proteins CSA (*ERCC8*) or CSB (*ERCC6*) sensitizes human cancer cells to the ICL inducing drug oxaliplatin^16^, suggesting that those proteins are required for repair of ICLs.

Indeed, CSB was found to promote repair of ICLs during transcription, in conjunction with TLS polymerase Pol ζ^17^.

CSA and CSB play a critical role in transcription-coupled nucleotide excision repair (TC-NER), leading to the hypothesis that TC-NER could process ICLs. However, this model has not been fully established. TC-NER is triggered by lesions that halt transcribing RNA Polymerase II (Pol II), such as ultra violet (UV) induced pyrimidine dimers or bulky DNA adducts^18^. Pol II stalled at the lesion recruits TC-NER factors CSA, CSB, USP7, and UVSSA to remodel the transcription complex by eviction of DSIF and RFT1 as well as ubiquitylation of subunit RPB1^19,20^. The modified Pol II complex backtracks along the DNA, exposing the lesion for repair. TFIIH is then recruited and promotes dual endonuclease cleavage, removing the damaged DNA segment from the transcribed strand. Gap filling synthesis and subsequent ligation restores the transcribed strand^21,22^.

The UVSSA protein functions in TC-NER, and mutations in the *UVSSA* gene cause the UV sensitivity syndrome (UVSS)^23–26^. UVSSA is recruited to stalled Pol II by CSA, and in turn facilitates the recruitment of the USP7 deubiquitinase and TFIIH to stalled Pol II. Loss of UVSSA impairs TC-NER and sensitizes cells to UV damage^27,28^. Additionally, UVSSA preserves genome stability during oncogene-driven transcriptional stress, as it is needed for cell survival of MYC-dependent hyperactive transcription^29^. Collectively, UVSSA plays an important role in regulation of transcription during genomic stress.

UVSSA was independently linked to cell survival following ICL damage in three distinct genome-wide CRISPR knockout screens, indicating that loss of UVSSA sensitized cells to the ICL inducing drugs maphosphamide^30^ (table s1) and cisplatin^31,32^. Given UVSSA’s role in transcription-associated genome maintenance, these results suggest that UVSSA functions in TC-ICR.

Here we show that UVSSA promotes TC-ICR. UVSSA knockout sensitizes cells to crosslinking drugs. Cell survival following ICL damage is mediated by UVSSA binding to TFIIH. Repair of crosslinker induced DNA damage is significantly delayed in UVSSA^-/-^ cells. Using an ICL repair reporter assay, we show that UVSSA is required for transcription-coupled repair of a single ICL. Finally, we characterize the UVSSA interactome following ICL damage and document the overlap between UV and ICL dependent UVSSA interactions. Together, our results reveal UVSSA as a key ICL repair component during transcription, and identify a common role for UVSSA in TC-NER and TC-ICR.

## RESULTS

### UVSSA loss sensitizes cells to DNA crosslinking drugs and delays crosslinker damage repair

UVSSA was a significant result in multiple genome-wide CRISPR knockout screens for genes that regulate crosslinker sensitivity^30–32^, including a screen in acute lymphocytic leukemia (ALL) cells, searching for potential regulators of maphosphamide sensitivity^30^ (Table S1). Loss of UVSSA resulted in enhanced crosslinking drug sensitivity, suggesting that UVSSA is required for ICL repair, possibly in a transcription-coupled mechanism. We sought to validate these findings in UVSSA knockout haploid Hap-1 cells (Fig S1A) and a UVSSA knockout Hap-1 cell line stably expressing a FLAG-UVSSA construct (Fig S1B). We compared the sensitivity of Hap-1 wild type (WT), UVSSA knockout (UVSSA^-^) and UVSSA^-^ cells expressing FLAG-UVSSA (UVSSA^-^ + FLAG-UVSSA) to cisplatin in a clonogenic assay. We showed that UVSSA^-^ cells are sensitive to cisplatin and that the sensitivity was due to a lack of UVSSA, as it was restored to wildtype levels in cells expressing FLAG-UVSSA (Fig 1A). UVSSA^-^ cells are also sensitive to the crosslink inducing drug mitomycin C (MMC) (fig S1C). These results establish a role for UVSSA in promoting cellular survival following exposure to ICL inducing drugs.

**Fig 1.**
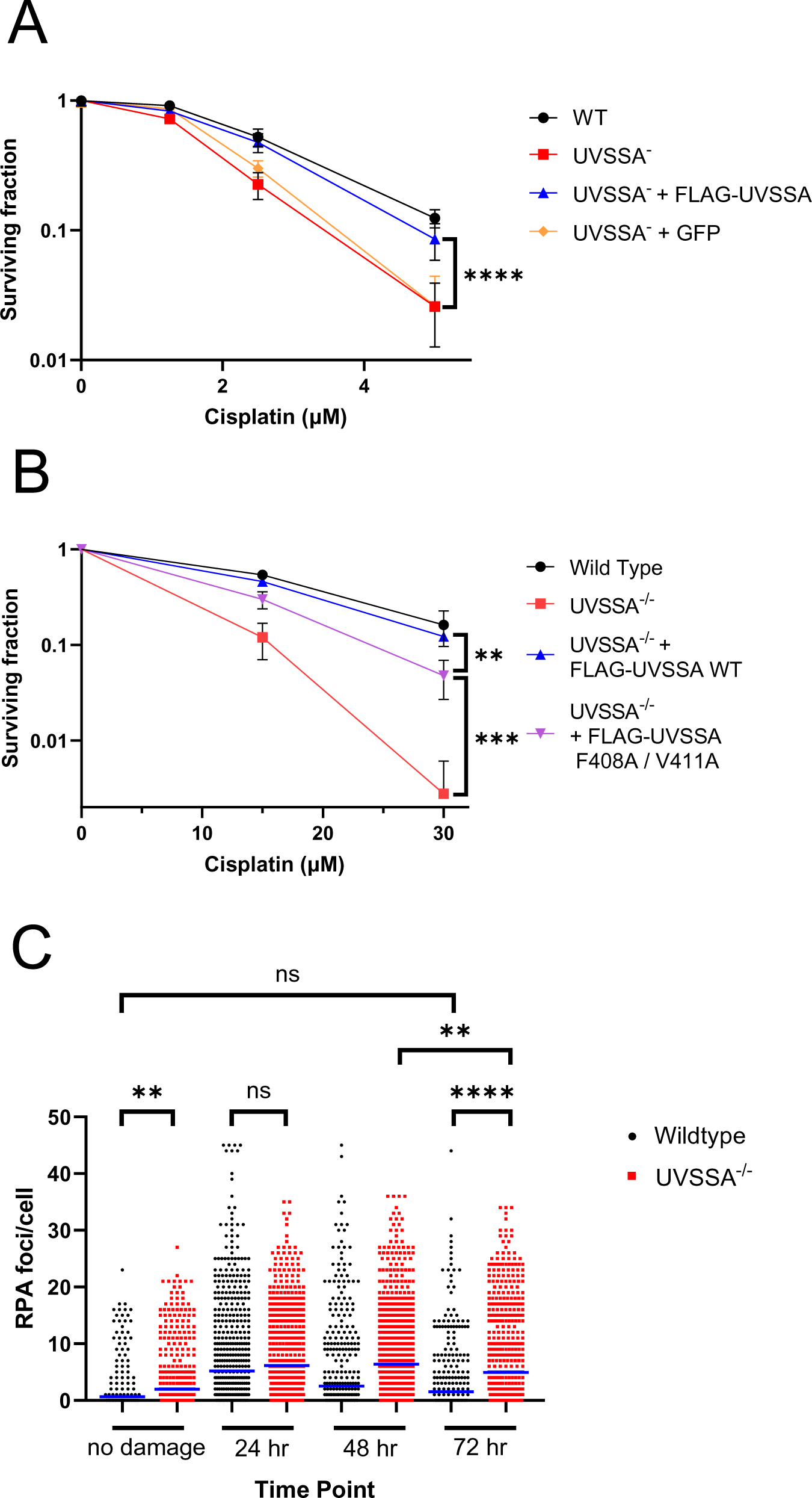
U**V**SSA **deficient cells are sensitive to cisplatin A)** Clonogenic assays monitoring WT, UVSSA^-^, UVSSA^-^ expressing GFP, and UVSSA^-^ expressing FLAG- UVSSA WT Hap-1 cell survival after 4-hour cisplatin exposure at the indicated doses. **B)** Same as in (A) for WT, UVSSA^-/-^, UVSSA^-/-^ expressing FLAG-UVSSA WT, and UVSSA^-/-^ expressing FLAG-UVSSA F408A/ V411A MCF10a cells. For A and B, N=6, data is mean and standard deviation of biological replicates. Statistical analysis by two tailed *t-*test, ** *p* <0.01, *** *p* <.001, **** *p* <0.0001. **C)** Quantification of RPA foci per cell in WT and UVSSA^-/-^ MCF10a cells at the indicated times following treatment with 10nM SJG-136 for 1 hour. Mean is indicated by a solid blue line. For each column left to right mean = 0.63, 1.79, 5.19, 6.15, 2.54, 6.38, 1.52, 4.95 N= 757, 734, 767, 690, 779, 772, 772, 743. Statistical analysis by one way ANOVA with multiple comparisons, \*\**p* <0.01, **** *p* <0.0001

We also generated a UVSSA^-/-^ line in diploid MCF10a cells, a non-transformed epithelial breast cell line, via CRISPR based iSTOP mutagenesis^33^. We expressed a nuclease-dead Cas9 fused to APOBEC1 along with a guide RNA to generate a W347* mutation in the *UVSSA* sequence (Methods). Mutation was confirmed by RFLP analysis and sequencing, and loss of expression was confirmed by western blot (Fig S1A). As anticipated, UVSSA^-/-^ MCF10a cells are also sensitive to cisplatin (Fig 1B).

UVSSA plays a role in TC-NER to repair UV damage^23,34^. In this pathway, UVSSA recruits repair factor TFIIH via interaction with UVSSA’s residues F408 and V411. Mutation in these residues blocks TFIIH binding and impairs transcription-coupled repair^28^. We sought to assess whether the UVSSA-TFIIH interactions were similarly required for UVSSA dependent ICL repair. We generated a FLAG-UVSSA F408A/ V411A coding sequence (Fig S1D), and isolated MCF10a UVSSA^-/-^ cell lines expressing the exogenous wild type and mutant FLAG-UVSSA protein at similar levels (Fig S1E). Expression of TFIIH binding deficient UVSSA only partially rescues cisplatin sensitivity. This suggests that UVSSA-TFIIH interaction is required for survival of crosslinking drugs (Fig 1B).

DNA lesions, including ICLs, can be visualized by immunofluorescence microscopy as foci of repair proteins on chromatin. The appearance and disappearance of these foci correlates with initiation and completion of repair of DNA lesions, respectively. We chose to monitor Replication Protein A (RPA) foci formation following ICL induction. RPA is a repair protein that binds to single strand DNA (ssDNA) during numerous repair processes^35^. In particular, RPA foci form at sites of ICL damage and RPA is involved in replication independent ICL repair^36,37^. Thus, RPA foci formation and resolution can serve as an indicator of ICL repair kinetics.

Using immunofluorescence microscopy, we quantified RPA foci in wildtype and UVSSA^-/-^ MCF10a cells at 24-hour intervals following treatment with the ICL inducing drug SJG-136 (fig S1F, 1C). Of note, UVSSA^-/-^ cells displayed more RPA foci in untreated conditions, suggesting that UVSSA-deficient cells have elevated endogenous DNA damage. Nonetheless, SJG-136 treatment increased RPA foci counts to a similar extent in both genotypes, to a mean of 5.19 and 6.15 foci per cell 24 hours after damage in wildtype or UVSSA^-/-^ cells, respectively. This indicates that lack of UVSSA does not impact the ability of SJG-136 to generate DNA crosslinks. In wildtype cells, RPA foci counts declined over time to a mean of 2.54 at 48 hours and 1.52 at 72 hours. The mean foci count at 72 hours was similar to that of no damage control levels, indicating that the repair machinery of WT cells can resolve SJG- 136 induced DNA damage. In contrast, RPA foci in UVSSA^-/-^ cells only started to decline after 48 hours and remained significantly elevated at 72 hours, with a mean of 4.59. While ICLs are eventually repaired in UVSSA^-/-^ cells, as indicated by a decline in RPA foci between 48 and 72 hours, repair is delayed compared to WT cells. This finding strengthens the idea that UVSSA is required for repair of ICLs, and that deficient ICL repair contributes to the ICL sensitivity observed in UVSSA knockout cells.

SJG-136 induces both ICLs and other types of lesions, including intra-strand DNA crosslinks, that could also form RPA foci. We next designed an assay to specifically report ICL repair in the absence of any additional lesions.

### UVSSA functions in transcription-coupled ICL repair

We designed a reporter plasmid-based assay to specifically assess transcription-coupled repair of a single ICL lesion. The plasmid harbors an ICL-containing oligonucleotide inserted between a CMV promoter and the mEmerald coding sequence. The ICL is induced *in vitro* by the drug SJG-136^38^, and the single lesion bearing oligonucleotide is purified to ensure that no other lesion types are present prior to ligation (see Methods). Upon transfection of the reporter plasmid into cells, the ICL blocks expression of the mEmerald reporter unless the lesion is repaired. mEmerald associated fluorescence is quantified to calculate repair efficiency (Fig 2A and Methods). Similar assays have previously been used to measure replication independent ICL repair^11,17,39^. The reporter plasmid lacks an origin of replication; thus, the readout is independent of replication and specific to transcription-coupled repair.

**Fig 2.**
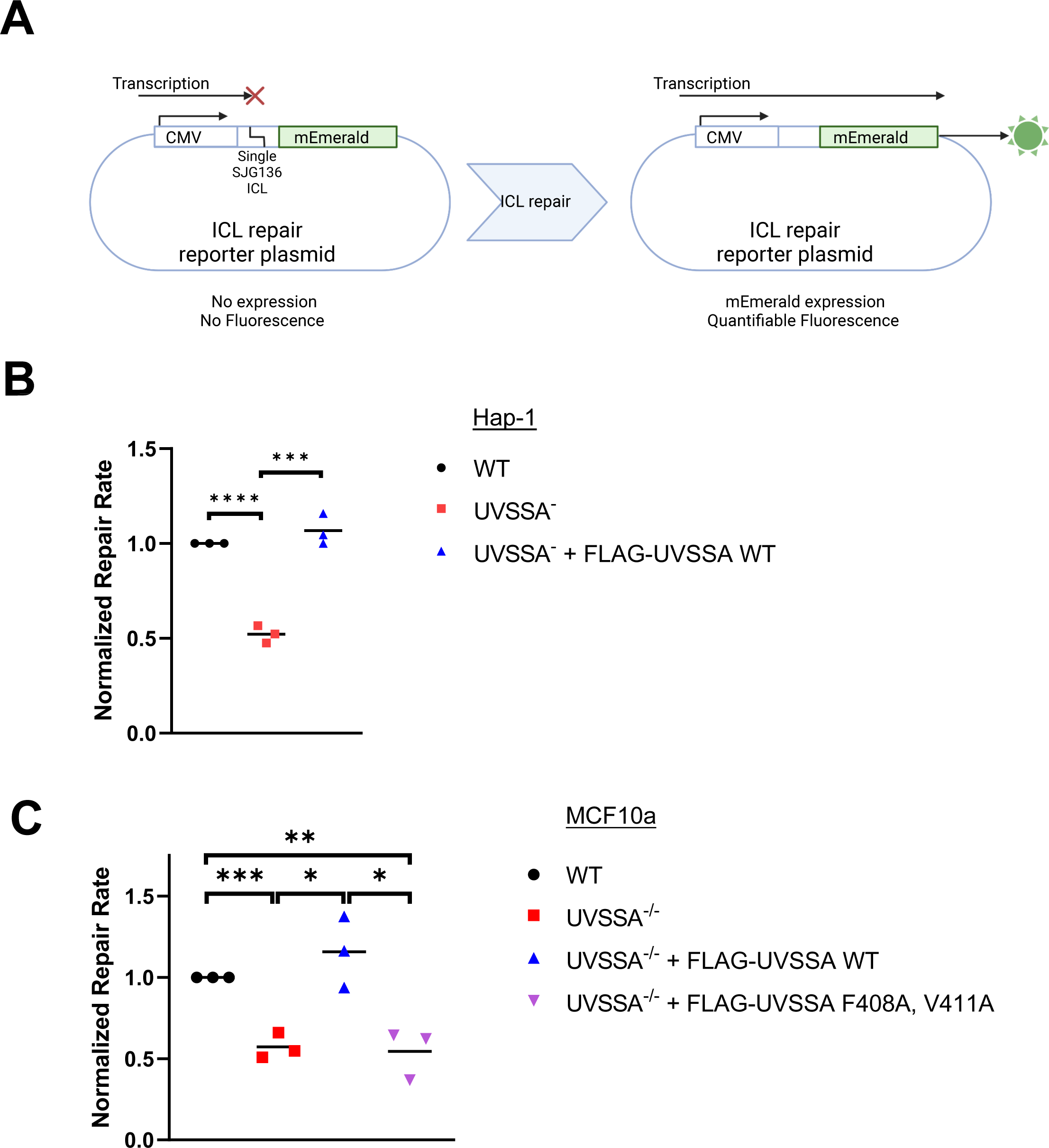
UVSSA is required for transcription-coupled repair of a single ICL **A)** Schematic of the reporter plasmid to monitor transcription-coupled ICL repair. A crosslink bearing oligonucleotide is inserted between the CMV promoter and an *mEmerald* coding region. Upon transfection, the ICL prevents *mEmerald* until it is repaired. (Created using Biorender.com). **B)** Normalized ICL repair efficiency in WT, UVSSA^-^ and UVSSA^-^ expressing FLAG-UVSSA Hap-1 lines transfected with the ICL reporter. **C)** Normalized ICL repair efficiency in WT, UVSSA^-/-^, UVSSA^-/-^ expressing FLAG-UVSSA WT and UVSSA^-/-^ expressing FLAG-UVSSA F408A/ V411A MCF10a lines transfected with the ICL reporter. For B and C, N=3 independent experiments, mean is indicated by a solid line. Repair rates are normalized to WT for each independent experiment. Statistical analysis by two tailed *t-*test, * *p* <0.05, ** *p* <0.01, *** *p* <0.001, **** *p* <0.0001

We transfected Hap-1 WT, UVSSA^-^, and UVSSA^-^ cells expressing FLAG-UVSSA with the reporter. UVSSA loss reduced ICL repair efficiency by 50% in UVSSA^-^ cells. Expression of FLAG-UVSSA in UVSSA^-^ cells restored repair efficiency to WT levels (Fig 2B), confirming that the phenotype was caused by loss of UVSSA. Our results establish that UVSSA participates in TC-ICR.

To test the role of the UVSSA-TFIIH interaction in ICL repair, we transfected MCF10a WT, UVSSA^-/-^, and UVSSA^-/-^ cells stably expressing WT or FLAG-UVSSA F408A/V411A with the ICL repair reporter plasmid. While WT FLAG-UVSSA was able to rescue the UVSSA knockout associated repair defect, TFIIH binding deficient UVSSA was not, confirming that the UVSSA-TFIIH interaction is required for TC-ICR (Fig 2C).

Our results indicate that UVSSA is required for repair of an ICL lesion that blocks transcription and strengthen the idea that a replication independent, transcription-coupled ICL repair involving UVSSA is required for cell survival of crosslinking drugs. We next sought to characterize the mechanisms of UVSSA mediated TC-ICR by analyzing UVSSA localization and protein interactions induced by ICL damage.

### UVSSA interacts with transcribing Pol II and transcription-coupled repair factors during TC-ICR

We hypothesized that UVSSA should localize to ICL-induced lesions and be enriched on chromatin upon ICL damage. Thus, we fractionated UVSSA^-^ Hap-1 cells expressing FLAG-UVSSA treated with SJG-136 or vehicle. We then probed chromatin and soluble fractions for FLAG-UVSSA (see Methods) by western blot, and found that FLAG-UVSSA was enriched in the chromatin fraction upon SJG-136 treatment (Fig 3A, quantified Fig S2a).

**Fig 3.**
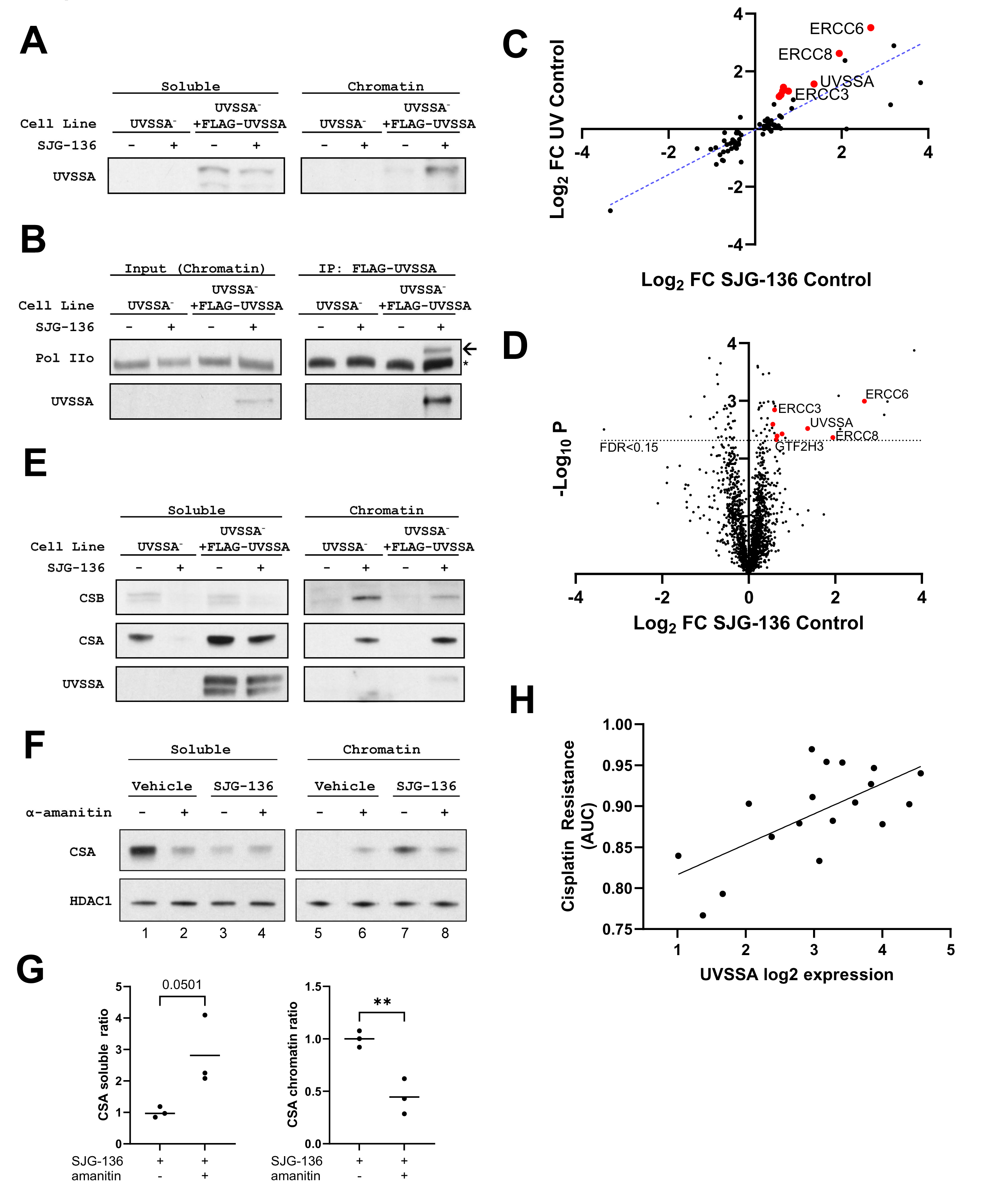
UVSSA localizes to ICL-damaged chromatin and interacts with transcription and repair proteins upon ICL damage **A)** Cell fractionation into soluble and chromatin fractions of UVSSA^-^ and UVSSA^-^ expressing FLAG- UVSSA Hap-1 cells treated with 100nM SJG-136 for two hours. **B)** Input and FLAG immunoprecipitation from chromatin fractions of UVSSA^-^ and UVSSA^-^ expressing FLAG-UVSSA Hap-1 cells treated with 100nM SJG-136 for two hours. Blots were probed with UVSSA antibody and an antibody recognizing the elongating form of Pol II: Pol IIo. Arrow indicates a Pol II band that specifically interacts with UVSSA upon DNA damage, * indicates a nonspecific band. This figure represents a composite of multiple representative western blots. **C)** Comparison of FLAG-UVSSA interactomes in UV or SJG-136 damaged hap-1 cells. Chromatin fractions of FLAG-UVSSA expressing Hap-1 cells treated with 100nM SJG-136 for 2 hours or exposed to 20 J/m^2^ UV-C were subjected to FLAG immunoprecipitation followed by mass spectrometry (see Methods). Proteins significantly enriched (FDR <0.15) are displayed. X-axis: Log2 fold change (log2 FC) following SJG-136 treatment, Y- axis : log2 FC following UV exposure. The dashed line is a linear regression, P<0.0001, N=74. Proteins with previously characterized repair function are highlighted in red. **D)** Volcano plot showing protein enrichment in FLAG-UVSSA pulldown from Hap-1 cell chromatin fractions following 100nM SJG-136 treatment for 2 hours. X-axis: Log2 FC upon SJG-136 treatment, Y-axis: *p* value of enrichment (two tailed t-test). **E)** Cell fractionation as in (A). Western blot is probed with the indicated antibodies. This figure represents a composite of multiple representative western blots. **F)** Cell fractionation of FLAG-UVSSA expressing UVSSA^-^ cells treated with 5ug/ml α-amanitin or vehicle 16 hours prior addition of 100nM SJG-136. Cell fractions were prepared after 2 hours of SJG-136 exposure. **G)** Quantification of normalized CSA soluble: total (left) or chromatin bound: total (right) ratios from Fig S2I (Methods). Analyzed by two tailed *t*-test, ** *p* < .01, N=3. Mean indicated by solid line. **H)** Correlation between UVSSA expression (RNA-seq), X axis and cisplatin resistance (Area Under the Curve, Y axis in Acute Lymphocytic Leukemia lines from CCLE database (methods). Linear regression analysis is depicted by a solid line, *p* = 0.0027, N=18

We next examined UVSSA interaction with RNA polymerase (Pol II) via co-immunoprecipitation (CoIP). UVSSA^-^ Hap-1 cells expressing FLAG-UVSSA were treated with SJG-136, fractionated, and anti-FLAG M2 affinity gel was used to isolate FLAG-UVSSA-bound proteins in chromatin fractions. The presence of Pol II was probed using an antibody detecting the RPB-1 subunit phosphorylated at serine 2, which is specific to elongating Pol II: termed Pol IIo^40^. SJG-136 treatment triggered the appearance of a Pol IIo band, indicating that UVSSA interacts with transcribing Pol II upon ICL damage (Fig 3B, quantified fig S2B). UVSSA-Pol IIo interaction was specific to the chromatin fraction (Fig S2C). UVSSA-Pol IIo interaction was also induced by another crosslinking drug, MMC, and by UV irradiation (Fig S2D), as previously described for TC-NER^34^. Our results strongly suggest that UVSSA interacts with transcribing Pol II to facilitate TC-ICR.

These data also point to analogous roles for UVSSA in transcription-coupled UV and ICL damage repair, and further supports the conclusion that TC-ICR is distinct from replication-coupled ICL repair. Therefore, we sought to compare the UVSSA interactomes following SJG-136 treatment or UV irradiation. We immunoprecipitated FLAG-UVSSA from chromatin fractions of mock treated, SJG-136 treated, or UV irradiated Hap-1 UVSSA^-^ cells expressing FLAG-UVSSA, followed by mass spectrometry (MS). We find that UVSSA interacts with a large set of overlapping partners following ICL or UV damage. Comparison of enrichment values upon UV or SJG-136 damage reveals a significant positive correlation between damage-induced UVSSA interacting proteins (fig 3C). These findings reveal a functional similarity between UVSSA’s role in repair of ICLs or UV photolesions, indicating an involvement of TC-NER processes in TC-ICR.

Examination of specific UVSSA interactions triggered upon SJG-136 treatment provides further evidence to support this conclusion. SJG-136 treatment induced interactions between UVSSA and transcription-coupled repair proteins CSA (*ERCC8*) and CSB (*ERCC6*), as well as several TFIIH components (*GTF2H4, ERCC3*, etc) (Fig 3D, Table S2), suggesting that CSA and CSB also play a role in ICL repair. Indeed, CSA and CSB are the 7^th^ and 4^th^ most enriched proteins, respectively, following SJG-136 treatment (Table S2). To validate this observation, we examined CSA and CSB localization in SJG-136 treated Hap-1 cells. We observed that both proteins are enriched on chromatin upon ICL damage. Chromatin localization occurs in both UVSSA^-^ and FLAG-UVSSA expressing cells (fig 3E, quantified Fig S2E,F), indicating that CSA and CSB localize to chromatin independently of UVSSA. We observed FLAG-UVSSA interaction with CSA upon ICL damage by Co-IP, (Fig S2G) confirming our MS results.

UVSSA interacts with transcribing Pol II in ICL damaged cells. Thus, we sought to assess whether localization of UVSSA, CSA, and CSB proteins to ICL damaged chromatin was dependent on ongoing transcription and sensitive to transcription inhibitors. α-amanitin binds to the RPB-1 subunit of Pol II and prevents translocation of DNA and RNA through the active site, therefore blocking elongation^41^. If Pol II is functioning as the damage sensor in TC-ICR, then α-amanitin treatment should impair the localization of TC-ICR proteins to chromatin by preventing Pol II dependent ICL detection and TC-ICR activation.

We monitored CSA protein localization in SJG-136 treated cells in the presence of 5μg/ml α- amanitin, a dose that did not significantly impact cell viability. SJG-136 induced CSA chromatin localization decreased in α-amanitin treated cells (Fig 3F, lanes 7 and 8). However, α-amanitin and SJG-136 treatments both impact CSA steady state levels (Fig S2H). To account for this decrease in total CSA protein, we calculated the ratio of soluble and chromatin bound CSA to total CSA (see Methods). Upon normalization, we observe a significant decrease in SJG-136 induced chromatin bound CSA in α-amanitin treated cells (Fig 3G right, Fig S2I). Conversely, soluble CSA increases in α- amanitin treated cells (Fig 3G left, S2I), indicating that a greater fraction of soluble CSA remains unbound to the chromatin. These results indicate that α-amanitin treatment prevents CSA localization to ICL damaged chromatin. We conclude that transcription inhibition via α-amanitin prevents CSA assembly into the TC-ICR complex. These findings support our hypothesis that TC-ICR is transcription dependent.

Our findings suggest that TC-ICR may facilitate ICL repair in low proliferating cells and promote resistance to crosslinkers cancer therapies^9^. Overexpression of repair factors has previously been linked to chemoresistance^42,43^, as cancer cells engage repair pathways to prevent DNA damage induced genotoxicity. We compared UVSSA expression and crosslinker drug resistance of cancer cell lines from the Cancer Cell Line Encyclopedia using DepMap cancer dependency database (depmap.org). We observed a significant positive correlation between UVSSA expression levels and cisplatin resistance in several cancer cell types, including acute lymphocytic leukemia (fig 3H), breast ductal carcinoma, and esophageal lineage cancers. A similar correlation was observed ween MMC resistance and UVSSA expression in the same cell types (Table 1). Our results suggest that chemoresistance to crosslinking drugs could arise in part via UVSSA-dependent activation of the TC- ICR pathway.

**Table 1.** Crosslinking chemotherapy resistance and UVSSA expression levels are correlated in several cancer types. Table of cancer types displaying a significant correlation between crosslinking drug resistance and UVSSA expression in the CCLE database (Methods). Slope, Pearson correlation, and *p* values were generated by linear regression analysis using the depmap web portal (https://depmap.org/portal/)

| Cell Type | Drug | Slope | Pearson | P Linear Regression |
| --- | --- | --- | --- | --- |
| ALL | Cisplatin | 12.5 | 0.701 | 9.2e-4 |
| ALL | Mitomycin C | 3.3 | 0.491 | 5.35e-2 |
| Esophageal | Cisplatin | 7.66 | 0.399 | 3.91e-2 |
| Breast ductal carcinoma | Cisplatin | 12.2 | 0.62 | 4.65e-3 |
| Breast ductal carcinoma | Mitomycin C | 1.9 | 4.53 | 2.61e-2 |

## DISCUSSION

ICLs are cytotoxic lesions that arise as consequences of endogenous metabolism or are induced by chemotherapeutic drugs. ICLs are highly genotoxic as they impede transcription and replication^1–3^, and mutagenic if improperly repaired. Replication-coupled repair of ICLs requires the FA pathway^4,5^. Replication independent repair (RIR) of ICLs may engage the Mismatch Repair (MMR) pathway^12^, Global Genome-Nucleotide Excision Repair (GG-NER)^13^ or Transcription-Coupled ICL Repair (TC-ICR) as reported in the context of CSB^17^. Here we identify UVSSA as a key player in TC-ICR.

### UVSSA is required for transcription-coupled ICL repair

We establish that UVSSA participates in ICL repair. First, UVSSA loss sensitizes cells to ICL inducing drugs cisplatin and MMC. Second, UVSSA is required for timely repair of DNA lesions in cells treated with the crosslinking agent SJG-136. Third, UVSSA is required for ICL repair in a fluorescence-based reporter assay specifically reporting transcription-coupled repair. Finally, UVSSA expression correlates with crosslinker resistance in several cancer types.

TC-ICR is challenging to study given that in dividing cells replicative polymerases scan the entire genome during S-phase to activate replication-coupled ICL repair. TC-ICR studies thus require experimental systems which allow monitoring replication-independent ICL repair. Non-replicating cell free systems and *in vitro* reporter reactivation assays have helped circumvent this hurdle. Cell free extracts from *Xenopus laevis* eggs that do not undergo replication have been used to study RIR of ICLs^11^ and to characterize MutSα-mediated repair of ICLs via MMR machinery^12^. Plasmid reporter assays have been employed to study transcription-coupled ICL repair by monitoring repair of transcription blocking lesions. Insertion of a purified ICL bearing oligonucleotide between a strong promoter and a reporter gene allows measurement of transcription-coupled repair and bypass of ICLs^17,39^. We have designed a fluorescence-based reporter that lacks an origin of replication, ensuring that we specifically measure transcription-coupled ICL repair.

Another important consideration when studying ICL repair is that crosslinking drugs induce DNA inter- and intra-strand crosslinks as well as monoadducts which all trigger a DNA damage response. Of these lesions, ICLs are the most cytotoxic and are a major cause of crosslinking induced toxicity. However, study of ICL repair defects cannot rely exclusively on experiments using crosslinking drugs in cellular assays due to the diversity of lesions induced during exposure. The fluorescence-based reporter plasmid described here addresses this concern, as the ICL inserted into the backbone is isolated and purified to ensure that no other lesion is present in the reporter. As such, the ICL repair reporter assay is the most direct and unambiguous experiment available to us to assay transcription coupled ICL repair in replicating cells.

We find that UVSSA is required for repair of a single ICL, thus establishing UVSSA as critical for transcription-coupled repair of ICLs. In addition, we show that UVSSA localizes to damaged chromatin in cells treated with crosslinking drugs, where it interacts with transcribing Pol II. Moreover, UVSSA interacts with transcription-coupled repair proteins CSA, CSB, and TFIIH upon ICL damage. Notably, loss of UVSSA-TFIIH interaction impairs transcription-coupled ICL repair. Knowing that UVSSA is involved in ICL repair, we can infer that these protein-protein interactions are also a component of TC-ICR. Finally, we show that ICL induced recruitment of CSA to chromatin is dependent on transcription, validating the concept of transcription-coupled ICL repair. MS analysis of UVSSA-interacting proteins failed to identify FA proteins upon ICL damage, suggesting that TC-ICR is independent of replication-coupled ICL repair.

### UVSSA: a common node in NER and TC-ICR

From the proteomics analysis of SJG-136 induced UVSSA protein interactions, we can build a mechanistic model of TC-ICR. We propose that TC-ICR is initiated when transcribing Pol II encounters an ICL, stalls and recruits CSA and CSB. UVSSA then localizes to Pol II, in turn recruiting TFIIH to the lesion. This characterization of UVSSA’s function in TC-ICR supports the previous models in which CSB participates in ICL repair during transcription^17^. In support of this model, we show: 1) that UVSSA interacts with transcribing Pol II following either UV, MMC, or SJG-136 induced damage; 2) a high degree of overlap between UVSSA interactomes following UV or ICL damage; and 3) that TFIIH recruitment is required for repair of both UV and ICL lesions. Together, our results provide the strongest evidence to date that TC-ICR is initiated via TC-NER lesion detection steps. Further research is needed to unravel the mechanisms that couple lesion recognition to downstream lesion removal in TC-ICR. Evidence in the literature suggests that NER endonuclease and TLS processes could contribute to ICL repair following detection by CSA, CSB, and UVSSA.

Translesion synthesis (TLS) polymerases are required for RIR of ICLs, possibly contributing to lesion processing during TC-ICR. In yeast, NER machinery functions in conjunction with TLS polymerase Pol η to remove an MMC induced ICL^44^. Similarly, the TLS polymerase Pol κ, which functions in NER during the gap filling step^45^, is also required for RIR of ICLs in *Xenopus* cell free extracts^11^. Finally, the TLS polymerase Pol ζ has been directly implicated in TC-ICR in human cells^17^. It is likely that multiple TLS and replicative polymerases perform lesion bypass during TC-ICR, facilitated by PCNA as observed in RIR of ICLs^11^. Additionally, CSB recruitment of the exonuclease SNM1A to ICLs may facilitate lesion processing prior to TLS^46^, as observed in replication-coupled ICL repair^47^.

### TC-ICR is required for survival following ICL damage independently of replication-coupled repair

Cells lacking UVSSA are sensitized to crosslinking drugs and show a significant delay in ICL repair. Given the role of UVSSA in TC-ICR, this strongly suggests that TC-ICR is required for cell homeostasis in response to ICL damage. While replication-coupled ICL repair is likely functional in UVSSA deficient cells, we show that this pathway is not able to compensate for loss of TC-ICR. It may be that ICL repair by the FA pathway during replication is not sufficient to resolve all drug-induced crosslinks, which would suggest that replication-coupled and replication-independent ICL repair mechanisms are not redundant (Fig 4). Previous studies indicate that these pathways contribute independently to ICL resistance: combined inhibition of CSB and FA pathways results in additive sensitivity to crosslinking drugs^17^. The sensitivity of TC-ICR defective cells to ICLs could have multiple causes. ICLs forming within essential genes stall transcription, requiring rapid removal to resolve transcription blocks and restore homeostasis^48^. In the absence of TC-ICR, stalled RNA polymerases should accumulate, increasing the frequency of collision with DNA polymerases, resulting in Replication-Transcription Conflict (RTC). RTCs are mutagenic, can cause chromosome rearrangement^49^, and would occur regardless of FA pathway functionality. Alternatively, in the absence of UVSSA, TC-ICR could initiate but not complete repair, leaving a repair intermediate that cannot be processed by replication-coupled ICL repair. A similar situation has been described following the loss of Pol κ^11^.

**Figure 4.**
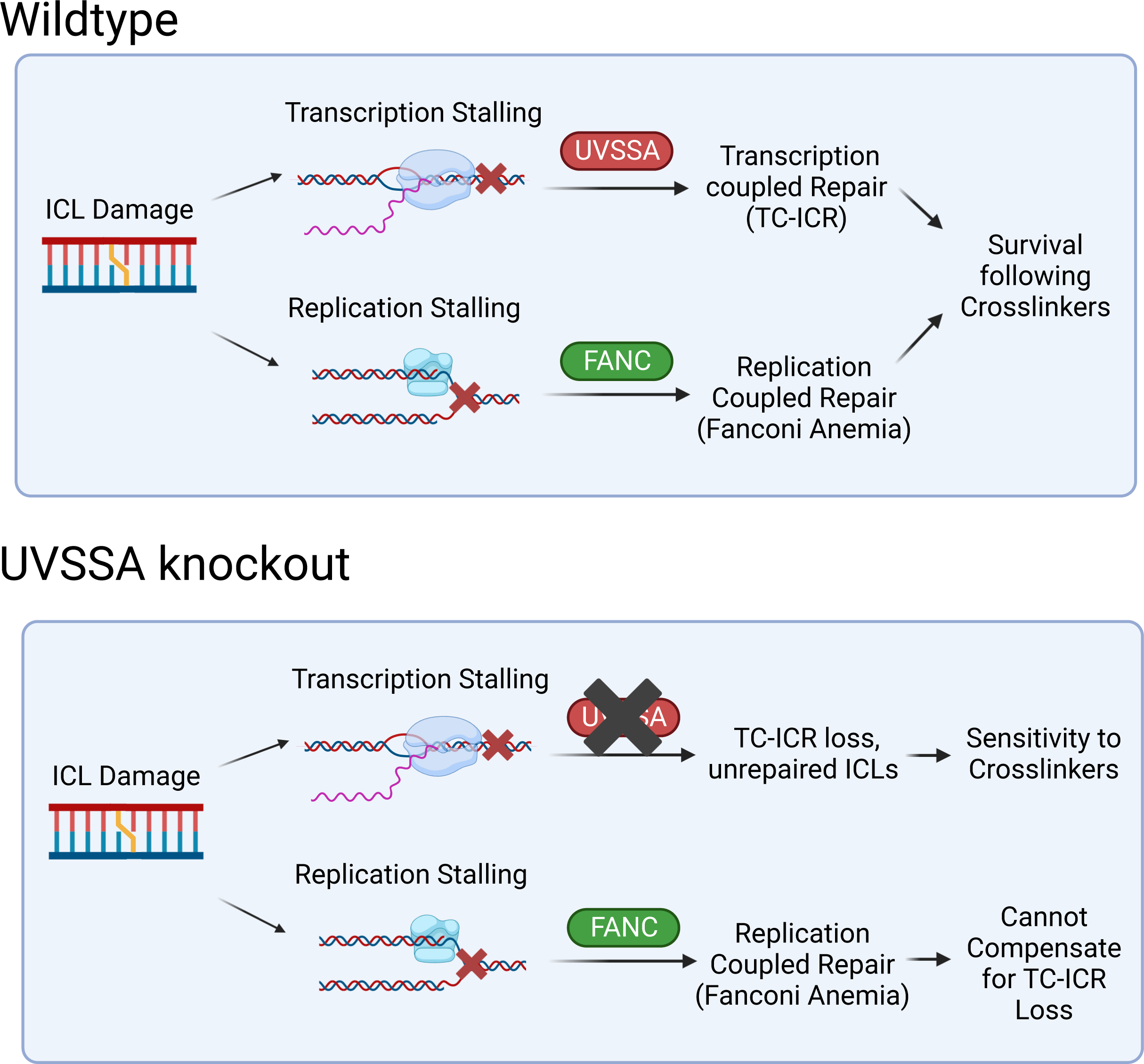
Contribution UVSSA to transcription coupled repair of ICLs Top: Contributions of replication- and transcription-coupled repair to survival from crosslinking drugs. UVSSA-dependent TC-NER and FANC-dependent pathways function independently to promote the survival to crosslinkers. Bottom: Lack of UVSSA-dependent TC-ICR is not compensated for by replication-coupled ICL repair. Created using Biorender.com

While loss of TC-ICR sensitizes cells, hyperactivation of the pathway may alleviate ICL induced transcription stalling and RTCs by increasing the efficiency of repair, promoting cellular resistance to crosslinking drugs. This could result in acquired chemoresistance, as observed in hyperactivation of other repair pathways in cancer^50^ and suggested by increased ICL resistance in high UVSSA expression cancer lines.

While TC-ICR is required for survival of crosslinker treatment, it might have a modest impact on ICL repair in unstressed growth conditions. Defects in the FA pathway, involved in replication-coupled ICL repair, result in bone marrow failure and cancer predisposition, highlighting the importance of ICL repair during normal replication^1^. In contrast, UVSS patients, defective in TC-NER and TC-ICR, display UV sensitivity without cancer predisposition or other developmental abnormalities^51^. CSA and CSB are also likely required for TC-ICR^17^ and are mutated in the TC-NER deficiency disease Cockayne syndrome (CS), characterized by developmental delay and short lifespan^52^. These patients are likely also defective in TC-ICR, but the divergence in clinical presentation between UVSS and CS suggests that TC-ICR deficiency does not contribute significantly to CS phenotypes. CSA and CSB requirements for oxidative lesion repair^53^ or Pol II transcription processivity^20^ have been posited to explain the differences in CS and UVSS symptoms. Therefore, loss of UVSSA more specifically inhibits TC-NER and TC-ICR without compromising other transcription or repair functions, and analysis of UVSS phenotypes is the most reliable way to interpret the significance of TC-ICR for unstressed growth.

In light of this, it appears that TC-ICR might play a modest role in maintaining genome stability during development and rapid cell growth, while FA is vital. Spontaneous ICLs are rare and transcription stalling caused by endogenous ICL damage in UVSSA deficient cells may be resolved by replication-coupled ICL repair. Alternatively, other replication independent ICL repair mechanisms, such as MMR and GG-NER, may compensate for TC-ICR loss at physiological levels of ICL damage.

The phenotypes reported in FA patients indicate that TC-ICR is unable to compensate for loss of replication-coupled ICL repair. This is not surprising, as TC-ICR only operates in the transcribed genome, leaving ICLs in non-transcribed regions unrepaired in FA deficient cells. These observations support a model in which TC-ICR is specifically required for transcription stability during times of high ICL damage, most notably following cancer chemotherapy.

Finally, TC-ICR may be critical in the maintenance of long-term transcription fidelity during aging. Endogenous DNA damage in cardiomyocytes necessitates NER proteins to prevent apoptosis signaling and cardiac failure^54,55^, which is linked to age related decline of cardiac function^56^. Aging is associated with profound transcription profile changes, connected to Pol II stalling at unrepaired DNA damage^57^. Notably, progeria and neurodegeneration have been associated with endogenous ICL forming aldehydes in DNA repair deficiency syndromes^58^. It is probable that ICLs induced by metabolic byproducts disrupt the transcriptome by blocking Pol II progression, contributing to age-related decline. TC-ICR should counteract this by repairing transcription blocking ICLs, particularly in non or rarely-cycling cells such as neurons where the FA pathway is not active. Further research into the TC-ICR pathway is necessary to fully understand its role in transcriptome integrity and organism longevity.

Research into TC-ICR has the potential to elucidate mechanisms of cancer reoccurrence. Quiescent cancer cells contribute to reoccurrence due to their reversible exit from the cell cycle, as they do not replicate damaged DNA^59,60^. However, these cells must repair DNA damage before cell cycle reentry via replication independent repair. Following crosslinker treatment, TC-ICR is likely required to repair ICLs in gene bodies and maintain normal protein expression. If quiescent tumor cells are dependent on TC-ICR to survive crosslinker treatment, it could provide a druggable target to sensitize these cells to crosslinking agents and decrease the risk of cancer reoccurrence.

Our results indicate that UVSSA and TC-ICR may also inform cancer treatment regimens via precision oncology. The activity of repair proteins is associated with cancer patient outcomes; high ERCC1 expression is linked to poor chemotherapeutic response^42,43^ while deleterious mutations in NER pathway proteins correlate with improved prognosis^61^. Our analysis shows that UVSSA expression correlates with resistance to crosslink inducing drugs in some cancer types, indicating that UVSSA expression may have value as a biomarker to inform chemotherapeutic choice during cancer treatment. In conclusion, our work establishes UVSSA as a factor in transcription-coupled repair of ICLs, elucidates the mechanisms through which transcription stalling initiates repair of ICLs, and reinforces the independence of TC-ICR from replication-coupled ICL repair mechanisms.

## DATA AVAILABILITY

The mass spectrometry proteomics data have been deposited to the ProteomeXchange Consortium via the PRIDE^62^ partner repository with the dataset identifier PXD041455.

All other data underlying this article is accessible in Dryad with DOI 10.5061/dryad.rfj6q57g8 All materials generated during the work described in this text are available upon request.

## AUTHOR CONTRIBUTIONS

R. C. Liebau conducted the majority of experiments. C. Waters conducted initial experiments in clonogenic sensitivity, engineering of the pEm-N1-CW reporter plasmid, initial experiments in ICL repair efficiency, and helped design the sgRNA for UVSSA knockout. A. Ahmed conducted immunofluorescence microscopy experiments and analyzed the data. R.K. Soni conducted liquid chromatography and mass spectrometry with initial data analysis. R. C. Liebau, C. Water, A Ahmed, J. Gautier designed the study, analyzed data, and wrote the paper

## FUNDING

This work was supported in part by National Institute of Health [grant numbers 1R35 CA197606, 1P01 CA174653 to JG]. Funding for open access charge by National Institute of Health [grant number 1P01 CA174653].

## CONFLICT OF INTEREST

The authors declare no competing financial interests.

## ACKNOWLEDGEMENTS

We thank Dr Ciccia (Columbia University) and lab members for providing materials and guidance to conduct iSTOP knockout. We thank Dr. Tanaka (Osaka University) for the gift of the FLAG-UVSSA construct. We thank the Proteomics and Macromolecular Crystallography Shared Resource, Herbert Irving Comprehensive Cancer Center (HICCC) at Columbia University (National Institute of Health grant number 5P30-CA013696).

## MATERIAL AND METHODS

### Cell culture

Cells were cultured at 37°C with 5% CO2. Hap-1 cells (Horizon C631) were cultured in IMDM (Thermo Fisher Scientific) supplemented with 10% FBS (Fisher Scientific) and 1% Penicillin/ Streptomycin (Invitrogen). MCF10a cells (ATCC CRL-10317) were cultured in DMEM/F12 media (Thermo Fisher Scientific) with 5% Horse serμ (Invitrogen), 1% Penicillin/Streptomycin, 20μg/ml EGF (Peprotech AF- 100-15), 500ug/ml Hydrocortisone (Sigma-Aldrich H0888-1G), 100μg/ml Cholera toxin (Sigma-Aldrich C8052), and 10mg/ml Insulin (Sigma-Aldrich I0516). 293T cells (ATCC CRL-3216) were cultured in DMEM (Thermo Fischer Scientific) supplemented with 10% FBS and 1% Penicillin/ Streptomycin

### Clonogenic assay

Cells were seeded at a density of 500 cells per 10cm dish, in triplicate for each condition. Cells were allowed to grow for 24 hours before addition of drugs at indicated doses. After a 4-hour treatment, the medium was removed by aspiration and replaced with fresh medium. Cells were incubated for 7- 14 days and stained upon sufficient colony growth. Colonies were stained by fixation in 100% methanol for 5 minutes, followed by incubation with 0.5% Crystal Violet for 5 minutes. Plates were rinsed and dried, and colonies were counted using the ICY software or manual counting

### Immunofluorescence microscopy

Wildtype and UVSSA^-/-^ MCF10a cells were cultured on 8-well chamber slides and subjected to 10 nM SJG-136 treatment (MedChem Express: HY-14573) or vehicle for 1 hour and incubated at 37 C for either 24, 48, or 72 hours. Cells were washed with PBS once and then pre-extracted with cold 2% PBS-Triton X-100 for 90 seconds. Cells were then washed with cold PBS for 1 minute and then fixed with 4% PFA (Electron Microscopy Sciences: 157-4) for 10 min. Cells were washed with room temperature PBS for 5 minutes before permeabilization in 0.1% PBS-Triton X-100 for 10 min. Cells were then washed with PBS three times for 5 minutes each before incubation with blocking buffer (3% BSA in PBS-Tween 20) at room temperature for 1 hour. Cells were then incubated overnight at 4°C with primary RPA antibody (Abcam: ab2175, 1/250) diluted in blocking buffer under a Hybrislip (Invitrogen: H-18202). Following primary antibody staining overnight, hybrislip was removed and cells were washed with PBS for 5 minutes, three times. Cells were then incubated with Alexa 488 conjugated goat anti-mouse IgG (Abcam: ab150113, 1/1,000) secondary antibody and DAPI stain (Invitrogen, 1/10,000) diluted in PBS for 1 hour. Following incubation, cells were washed with PBS three times for 5 minutes in the dark. Slides were prepared using Vectashield Mounting Medium (Vector Laboratories: H-1000-10) and then coverslipped.

Slides were analyzed under 40x magnification using a Zeiss Axio Imager Z2 microscope, equipped with a CoolCube1 camera (Carl Zeiss). MetaCyte software (version 3.10.6) was used to detect nuclei stained with DAPI and to perform automated foci quantification with customized classifiers. For each time point, a minimum of 300 cells were analyzed.

### Protein whole cell lysis preparation

For whole cell lysates, cells were lysed in RIPA lysis buffer (NaCl 150mM, NP-40 1%, Deoxycholate 0.5%, Sodium Dodecyl Sulfate 1%, Tris HCl pH 8 50mM) 30 minutes on ice followed by high-speed centrifugation and collection of the supernatant. Protein concentration was quantified using the Pierce BCA Protein Assay Kit (Thermo Scientific 23225). Lysates were mixed with equal volume 2x Laemmli buffer (Sodium Dodecyl Sulfate 4%, 2-mercaptoethanol 10%, glycerol 20%, Bromophenol Blue 0.004%, Tris HCl pH 8 125mM) and boiled for 5 minutes at 95°C before storage at -20°C

### Chromatin fractionation

Cells were fractionated as described in^25^. Briefly, cells were lysed in fractionation buffer (Tris HCl pH 7.5, KCl 100mM, Sucrose 300mM, MgCl2 2mM, Triton 0.1%, CaCl2 1mM, Dithiothreitol 1mM) on ice for 10 minutes, followed by centrifugation 5 min 3800 xG at 4°C. The supernatant was collected and saved as the soluble fraction. The pellet was washed in buffer, followed by digestion with Micrococcal Nuclease in buffer at 1000 units/ml for 30 minutes at room temperature. Digestion was halted by addition of ethylenediaminetetraacetic acid to a final concentration of 5mM. Samples were centrifuged 5 min 3800 xG at 4°C and the supernatant was collected. The pellet was washed with fractionation buffer and centrifuged again, and the supernatants were combined to generate the chromatin fraction. Protein concentration was assayed by BCA as above, and lysates were then boiled in equal volumes 2x Laemmli buffer before storage at -20°C

### Western blotting

Samples were run on precast Tris-Glycine 8% Novex gels (Thermo Fisher Scientific XP00080BOX) with GTS running buffer (Tris 25 mM, Glycine 190 mM, SDS 0.1%) or 4-12% Bis-Tris precast NuPAGE gels (Thermo Fisher Scientific NP0322BOX) with MOPS running buffer (Thermo Scientific NP0001). For whole cell lysates, 60-30 μg of protein was loaded. For cellular fractions, 60-30μg or 30-10μg of protein was loaded for the soluble or chromatin fractions, respectively. Proteins were transferred to PVDF membrane using the Iblot 2 system (Thermo Fisher Scientific IB21001). Membranes were then blocked for 1-2 hours in 5% powdered milk (Amresco, M203-10G-10PK) in PBST. Membranes were incubated overnight at 4°C with diluted antibodies as indicated. Membranes were then washed in PBST for 5 minutes 4X, followed by incubation with appropriate HRP conjugated secondary antibody at 1:10000-1:50000 dilution for 1 hour at room temperature. Membranes were again washed in PBST 4X 5 minutes each, followed by chemiluminescence either with Peirce ECL western Blotting Substrate (Thermo Fisher Scientific 32106), SuperSignal West Pico Plus (Thermo Fisher 34580), or SuperSignal West Dura extended Duration Substrate (Fisher Scientific 37071). Hyblot CI autoradiography film (Thomas Scientific 1159M38) was exposed to the membranes, fixed, and scanned on an Epson Perfection 37V scanner (Epson B11B207201)

### Co-Immunoprecipitation

Cellular fractions generated following chromatin fractionation protocol were pre cleared by mixing with Sepharose beads (Sigma-Aldrich 4B200) for 30 minutes at 4°C. After centrifugation, the supernatant was extracted using a 30-gauge needle, and then incubated with Anti-Flag M2 affinity gel (Sigma-Aldrich A2220) overnight at 4°C. Beads were washed 5x steps in fractionation buffer, then proteins were eluted in 2x SDS sample buffer (Tris HCl pH 6.8 125 mM, SDS 4%, Glycerol 20%, Bromophenol Blue 0.004%) by boiling at 95°C. 5-10ul of elution sample was ran for western blot.

### Western blot quantification

Quantification was performed using ImageJ software. The mean gray value for each band was evaluated and processed to correct for background signal. This value was then normalized to loading control signal. The band intensity was then normalized to the average of untreated samples, if detectable.

To quantify the localized: total protein ratio in cell fractions, protein band intensity in whole cell lysate (WCL), soluble, and chromatin fractions was measured following treatment. The protein signal in WCL was quantified and normalized to the untreated control. This value was used to normalize the signal in soluble or chromatin fractions from similarly treated cells. This ratio approximates the portion of total protein that is present in each cellular compartment.

### On-beads digestion for mass spectrometry

The Co-IP protocol above was followed until completion of the final wash step. Immunoprecipitated proteins on agarose beads were then washed five times with 200 μl of 100 mM Tris-pH 8.0. Proteins were reduced with 10 mM TCEP and alkylated with 11 mM iodoacetamide (IAA) that was quenched with 5 mM DTT. Protein digestion was processed by adding 1 μg of trypsin/Lys-C mix and incubated overnight at 37°C and 1400 rpm in a thermomixer. The next day, digested peptides were collected in a new microfuge tube and digestion was stopped by the addition of 1% TFA (final v/v), followed by centrifugation at 14,000 x g for 10 min at room temperature. Cleared digested peptides were desalted on an SDB-RPS Stage-Tip^63^, and dried in a speed-vac. Peptides were dissolved in 3% acetonitrile/0.1% formic acid.

### Liquid chromatography with tandem mass spectrometry (LC-MS/MS)

Peptides were separated within 80 min at a flow rate of 400 nl/min on a reversed-phase C18 column with an integrated CaptiveSpray Emitter (25 cm x 75µm, 1.6 µm, IonOpticks). Mobile phases A and B were with 0.1% formic acid in water and 0.1% formic acid in ACN. The fraction of B was linearly increased from 2 to 23% within 70 min, followed by an increase to 35% within 10 min and a further increase to 80% before re-equilibration. The timsTOF Pro was operated in PASEF mode^64^ with the following settings: Mass Range 100 to 1700m/z, 1/K0 Start 0.6 Vs/cm-2, End 1.6 Vs/cm-2, Ramp time 100ms, Lock Duty Cycle to 100%, Capillary Voltage 1600V, Dry Gas 3 l/min, Dry Temp 200°C, PASEF settings: 10 MS/MS Frames (1.16 seconds duty cycle), charge range 0-5, an active exclusion for 0.4 min, Target intensity 20000, Intensity threshold 2500, CID collision energy 59eV. A polygon filter was applied to the m/z and ion mobility plane to select features most likely representing peptide precursors rather than singly charged background ions.

### LC-MS/MS data analysis

Acquired PASEF raw files were analyzed using the MaxQuant environment V.2.1.3.0 and Andromeda for database searches at default settings with a few modifications^65^. The default is used for the first search tolerance and main search tolerance (20 ppm and 4.5 ppm, respectively). MaxQuant was set up to search with the reference human proteome database downloaded from UniProt. MaxQuant performed the search trypsin digestion with up to 2 missed cleavages. Peptide, site, and protein false discovery rates (FDR) were all set to 1% with a minimum of 1 peptide needed for identification; label-free quantitation (LFQ) was performed with a minimum ratio count of 1. The following modifications were used for protein identification and quantification: Carbamidomethylation of cysteine residues (+57.021 Da) was set as static modifications, while the oxidation of methionine residues (+15.995 Da), and deamidation (+0.984) on asparagine were set as a variable modification. Results obtained from MaxQuant, protein groups table was further used for data analysis.

### CoIP protein enrichment analysis

LFQ values were extracted from MaxQuant analysis and further analyzed using Perseus software. Standard transformation including removal of “only identified by site” results and transformation by log2 was performed. Proteins displaying partial detection in one condition (mix of zeros and values) were eliminated, such that only proteins that were consistently detected or not detected remained. Zeros were replaced by imputation from a normal distribution (width 0.3, downshift 1.8) using Perseus tools. Finally, significance of enrichment was analyzed using students *t*-test comparing mock treated samples to SJG-136 or UV damaged samples. FDR was calculated using Benjamini-Hochberg method. Log2 FC was calculated by the Persius program.

### DNA interstrand crosslinked oligonucleotide preparation

Single stranded insert oligonucleotides (see oligo table) were brought to 95°C for 5 minutes and allowed to cool and anneal for 2 hours. 100μg of annealed oligo was mixed with 2x SJG-136 buffer (Triethanolamine 50mM, EDTA 2mM) and SJG-136 was added to a final concentration of 100μM.

Oligos were incubated overnight at 37°C before ethanol precipitation. The oligo was then run on a 15% PAGE urea denaturing gel to separate crosslinked from un-crosslinked species. The heavier running crosslinked species was excised from the gel via UV shadowing and then eluted by crush and soak method and isopropanol precipitation. The crosslink bearing oligo and an uncrosslinked control was then phosphorylated by T4 polynucleotide kinase (New England Biolabs M0201S).

### pmEmerald-n1 modification for ICL reporter assay

The pmEmerald-n1 plasmid was mutagenized to disrupt the Blp1 site at 1358 and to remove the SV40 replication site using two rounds of Quikchange II site directed mutagenesis (Aligent Technologies 200523). The reporter was digested using Bam HI and Nhe I to insert a fragment from the pEGFP-N3-ΔSV40 plasmid^11^ containing two BbsI sites, such that digestion with BbsI would generate overhangs compatible with insertion of a single crosslink bearing oligonucleotide. The resulting plasmid was dubbed pEm-N1-CW. Editing was confirmed by sequencing.

### ICL reporter backbone preparation

pEm-n1-CW was digested by BbsI. The digest was run on an ethidium bromide agarose gel and the linearized fragment excised under brightfield light. The linear DNA backbone was then extracted from the gel via electroelution in D-tubes (EMD Millipore 71508-3), followed by phenol chloroform extraction and butanol concentration.

### ICL reporter ligation and purification

pEm-n1-CW backbone was ligated to either crosslinked insert or control insert by T4 DNA ligase for 72 hours at 4°C. After confirming ligation by agarose gel electrophoresis, both reporters were purified by phenol chloroform extraction and concentrated by butanol extraction. The crosslink bearing reporter was then digested overnight with BglII Ito remove any non-crosslinked species.

Both reporter plasmids were then purified via CsCl gradient to remove linear products and desalted and concentrated using an amicon ultra filter tube (EMD Millipore UFC503024)

### ICL reporter transfection

Cells were transfected using the neon transfection system. Following the neon transfection protocol, 3*10^5^ Hap-1cells or 2*10^5^ MCF10a cells were transfected with 1ug pCAGGS carrier DNA, 50ng pmCherry-c1 for transfection efficiency control, and 50ng of either crosslinked or uncrosslinked control reporter. Cells were grown for 24 hours and then collected by trypsinization and resuspended in PBS before fluorescence acquisition on an Attune NxT Flow Cytometer (Thermo Fischer A29002) and gated for green (530/30 nm) and red (620/15 nm) emission.

### ICL repair efficiency calculation

FCS7 express software was used to analyze FACS data files. After standard gating to remove doublets, fluorescence gates were set based on a negative control to determine the percentage of cells fluorescing. The mEmerald fluorescing cell population was then subdivided based on fluorescence intensity into low, medium, and bright fluorescing groups. The percentage of cells that fell into the bright intensity gate was then calculated, and normalized to transfection efficiency calculated from mCherry fluorescence. The normalized bright value of cells transfected with the crosslinked reporter was then divided by the same value for the cell transfected with the control reporter, generating a repair efficiency value. Any technical repeats were averaged. Repair efficiency was then normalized to wild type.

### UVSSA knockout by iSTOP

The iSTOP sgRNA was designed using the web tool provided by Dr. Ciccia’s lab (https://www.ciccialab-database.com/istop/<u>#/</u>). The sgRNA guide was ligated into the B52 sgRNA expression plasmid. A plasmid expressing the BE3 enzyme, a modified Cas9 enzyme, and containing a blasticidin resistance marker, as described in^33^, was generously provided by the Ciccia lab. The BE3 expression vector and the sgSTOP expression vector were cotransfected via JetPEI (Polyplus 101000053) and cells were incubated for 72 hours before selection with blasticidin. The selected population was expanded to generate an uncloned population that was subsequently cloned to isolate edited cells. Successful editing was confirmed by RFLP, sequencing, and western blot analysis.

### Single cell cloning

Uncloned populations were collected by trypsinization. The suspension was then diluted to a concentration of 4.8 cells per ml. 200ul of this suspension was distributed into each well of a 96 well plate. Cells were incubated for 2-3 weeks before identifying single colonies, which were expanded as needed. iSTOP editing was again confirmed by RFLP analysis, sequencing, and western blot for protein expression.

### UVSSA mutagenesis

The FLAG-UVSSA fusion protein expressing plasmid (pcDNA3.1 FLAG-UVSSA) was a generous gift from the Tanaka lab. Mutations in UVSSA functional residues were induced using the Q5 mutagenesis system. Primers were designed using the NEB base changer portal (https://nebasechanger.neb.com). Mutation was confirmed by sequencing. The mutated UVSSA coding region was then amplified using primers designed to add AscI and EcoRI restriction sites, allowing ligation into the pBabe-puro backbone for retroviral transduction.

### Retroviral transduction

293T cells were transfected with pBabe puro plasmid containing either a wild type or mutated FLAG- UVSSA construct, along with VSV-G and pUVMC packaging plasmids via Jetpei (VWR 89129-916).

293T cells were grown for 24 hours before replacing the media with target cell media. After 24 hours the viral media was collected and filtered through a 0.45 micron filter before being added to cells along with 1 ml of fresh media and polybrene to a final concentration of 10ug/ml. 72 hours after transduction cells were selected with puromycin at a concentration of 2ug/ml. Cells were grown for 48 hours under selection before removal of the antibiotic and expansion of the population for cloning.

### Cancer Cell Line Encyclopedia data analysis

Cancer Cell Line Encyclopedia (CCLE) data was accessed using the DepMap portal (https://depmap.org/portal/ccle/). The portal was used to compare UVSSA expression (RNA seq) and drug resistance (area under the curve) for cancer lines in the database. The portal was used to subset data based on cancer lineage or disease type and calculate a linear regression between UVSSA expression and drug resistance values. Linear regression data was extracted from the portal, and a full data set for UVSSA expression and cisplatin resistance in acute lymphocytic leukemia lines was downloaded for independent analysis. Data was accessed from the Depmap Portal, Expression 22Q2 and Sanger GDSC1 data sets

### Statistical Analyses

Statistical significance of RPA foci counts was calculated using one way ANOVA with multiple comparisons in Graphpad Prism 9 software. Data was gathered from 3 biological replicates with N>300 for each datapoint. All other statistical values presented in the figures were calculated by appropriate *t-*test (paired, unpaired, two tail). Clonogenic survival data was gathered from 6 biological replicates per condition, N=6. ICL repair efficiency data was gathered from 3 independent biological replicates, with any technical repeats averaged, N=3. Western blot band intensity quantification was gathered from 3 biological replicates per condition, N=3. For protein enrichment, significance was analyzed by Perseus software (https://maxquant.net/perseus/) by *t-*test with multiple comparison correction by Benjamini-Hochberg method. CoIP protein enrichment data was collected from 3 independent biological replicates. All other comparisons were analyzed using Graphpad Prism 9 software (https://www.graphpad.com/). Linear regression analysis of DepMap data (table1) was generated by the DepMap web portal (https://depmap.org/portal/), all other regressions were analyzed by Graphpad Prism 9 software. A confidence interval of 95% (P<0.05) was set for all comparisons

### Reagents

Primary Antibodies: UVSSA (Genetex GTX106751), Vinculin (Cell signaling Technology 4650S), FLAG M2 (EMD millipore F3165), Pol IIo (EMD millipore 04-1571), CSA (Abcam ab137033), CSB (Santa Cruz Biology sc-166042), Histone 3 (Cell Signaling Technology 9715), HDAC1 (Abcam ab109411), RPA (Abcam ab2175).

Restriction enzymes: all restriction enzymes acquired from NEB: BbsI (R0539S), BamHI (R0136S), NheI-(HF R3131S), BglII (R0144S), AscI (R0558S), EcoRI (R0101S)

Kits: Q5 mutagenesis kit NEB (E0554S)

Equipment: Attune NXT flow cytometer (Thermo fisher scientific A29002). Neon transfection system (Thermo Fischer MPK5000).

### Biological Resources

Human cell lines: Hap-1 (Horizon C631), UVSSA^-^ (Horizon HZGHC005817c010). MCF10a (ATCC CRL- 10317). 293T (ATCC CRL-3216).

Plasmid vectors: pmEmerald-n1 (Addgene 53976), pBabe-puro (Addgene 10668), pCaggs (BCCM LMBP 2453), pMcherry-c1 (Addgene 58476), B52 (Addgene 100708), VSV-g (Addgene 8454), pUVMC (Addgene 8449)

### Websites and programs

iSTOP sgRNA database https://www.ciccialab-database.com/istop/#/^33^. Depmap cancer dependency database https://depmap.org/portal/. NEB base changer mutagenesis PCR design tool https://nebasechanger.neb.com. Perseus protein quantification software, https://maxquant.net/perseus/^66^. FCS express fcs file explorer 7, https://denovosoftware.com/.

Graphpad prism 9, https://www.graphpad.com/. ICY bioimaging analysis https://icy.bioimageanalysis.org/. FIJI image processing software https://imagej.net/software/fiji/ ^67^

---

**Figure S1.**
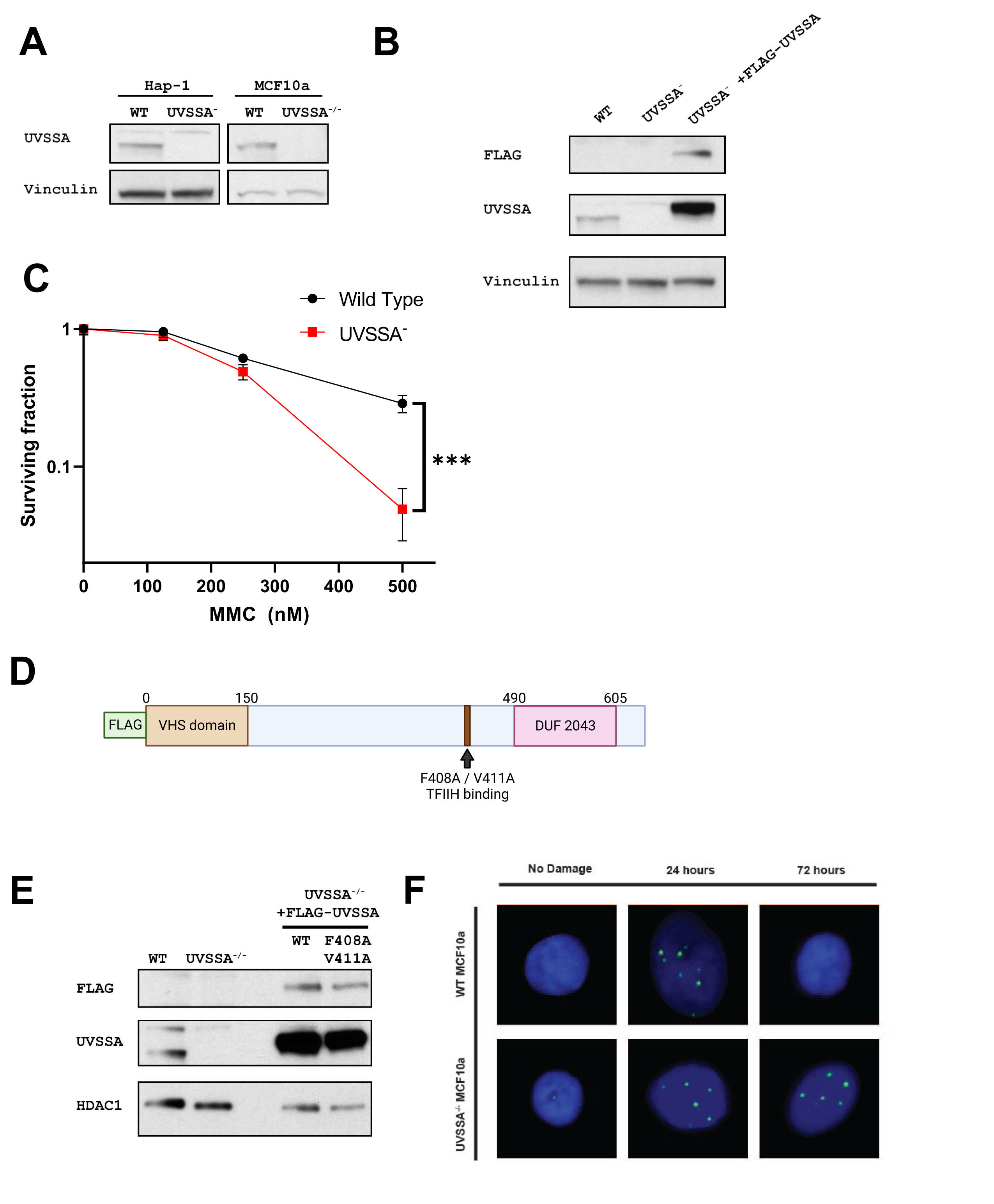
(Related to figure 1). UVSSA expression, MMC sensitivity **A)** UVSSA expression in Hap-1 WT or UVSSA^-^ and MCF10a WT or UVSSA^-/-^ cell. **B)** UVSSA expression in WT, UVSSA^-^, and UVSSA^-^ expressing FLAG-UVSSA Hap-1. **C)** Clonogenic assays monitoring WT and UVSSA^-^ Hap-1 cell survival following 4-hour Mitomycin C exposure at the indicated doses. N=6, data is mean and SD of biological replicates, statistical analysis by two tailed *t-*test, **** *p* < 0.0001. **D)** Schematic of UVSSA mutations abrogating TFIIH binding indicated. Created using Biorender.com. **E)** UVSSA expression in MCF10a cell lines expressing FLAG-UVSSA. **F)** Representative images of immunofluorescence microscopy monitoring RPA Foci at 24-hour intervals in WT and UVSSA^-/-^ MCF10a cells treated with 10nM SJG-136 for 1 hour. RPA foci green; nucleus/ DAPI: blue.

**Figure S2.**
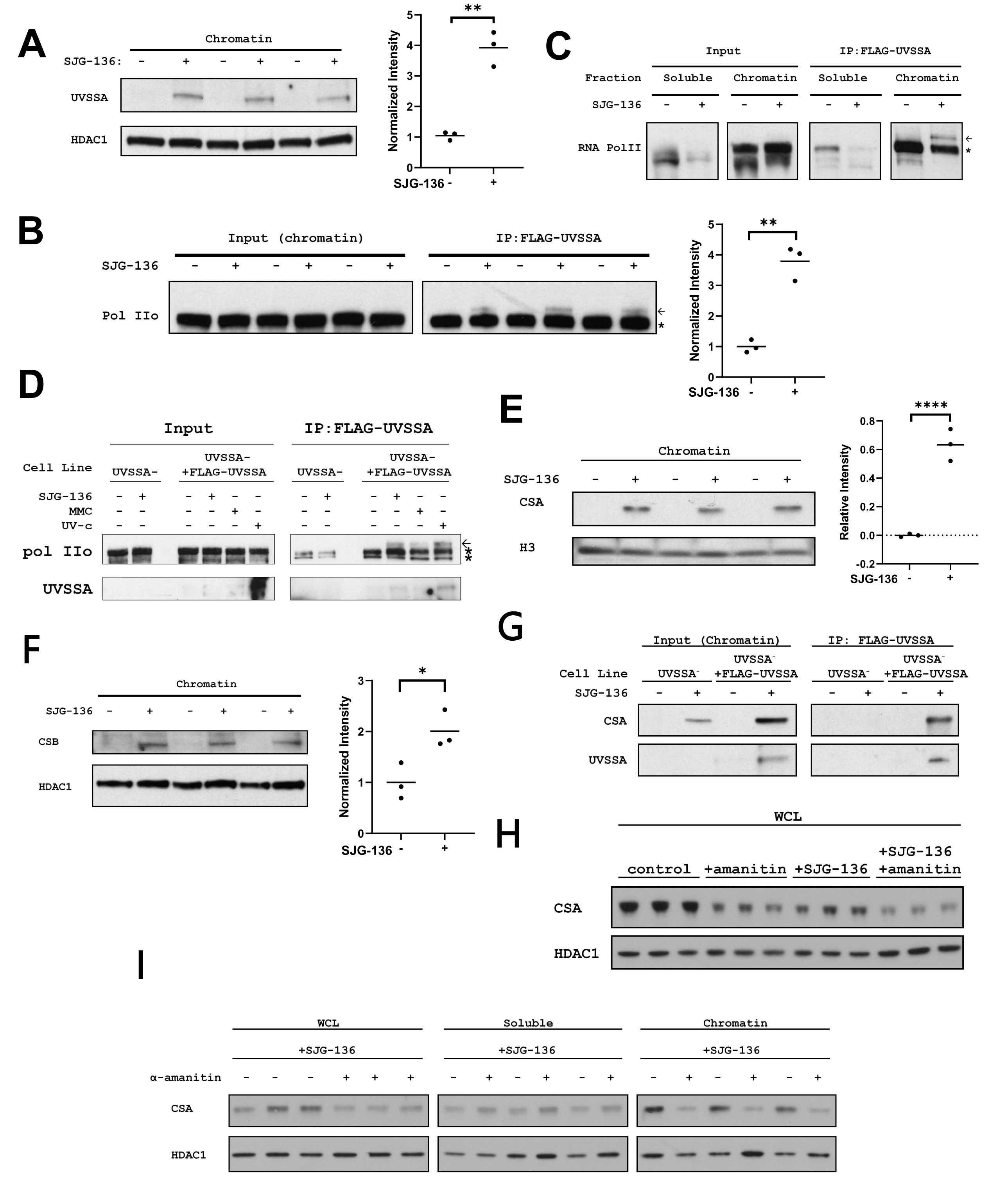
(related to figure 3) Quantification of protein localization and interaction upon ICL damage, expanded analysis of UVSSA-Pol II interaction **A)** Biological replicates of Fig 3A; chromatin bound UVSSA in FLAG-UVSSA Hap-1 cells with quantification. Each lane represents an independent biological experiment. HDAC1: loading control. **B)** Biological replicates of Fig 3B: Pol IIo pulldown and quantification in FLAG CoIP from UVSSA^-^ expressing FLAG-UVSSA Hap-1 cells. Each lane pair represents an independent biological repeat. Input signal was used as a loading control for analysis. **C)** Hap-1 cells treated with 100nM SJG-136 expressing FLAG-UVSSA were probed for Pol IIo in cell lysates (input) following FLAG CoIP of soluble and chromatin fractions. **D)** FLAG-UVSSA Hap-1 cells were treated with 100nM SJG-136 for 2 hours, 9μM MMC for 2 hours, or exposed to 20 J/m2 UV-C followed by FLAG IP. Western blot against Pol IIo in cell lysates (input) or FLAG CoIP is shown. For **B**, **C**, and **D**, arrow indicates a Pol II band that interacts with UVSSA upon DNA damage, * indicates a nonspecific band. **E, F)** Replicates of Fig 3E: chromatin-bound CSSA and CSB in FLAG-UVSSA Hap-1 cells with quantification. Each lane represents an independent biological repeat. Histone H3 or HDAC1 were used as loading controls for analysis. For panels **A**,**B**,**E**, and **F**, statistical analysis by two tailed *t-*test, * *p* <0.05, ** *p* <0.01, *** *p* <0.001, **** *p* <0.0001. The mean is indicated by a solid line. **G)** Input and FLAG immunoprecipitation from chromatin fractions of UVSSA^-^ and UVSSA^-^ expressing FLAG-UVSSA Hap-1 cells treated with 100nM SJG-136 for two hours. Western blots were probed with CSA and UVSSA antibodies. This figure represents a composite of multiple representative western blots. **H)** CSA western blot from UVSSA^-^ cells expressing FLAG-UVSSA construct treated with 5ug/ml α-amanitin for 16 hours followed by 100nM of SJG-136 for two hours. Each lane is an independent biological repeat. **I)** CSA western blots in WCL, soluble, or chromatin fractions from cells as in Fig 3F. Each lane is an independent biological replicate.

**Table S1.** UVSSA depletions sensitizes leukemia cells to Maphosphamide. Data from Oshima 2020^30^, subset of supplementary table 7g. Data from a CRISPR knockout screen in maphosphamide treated ALL cancer cells. Top hits with FDR threshold of 0.05 for guides significantly depleted in the screen are shown. Values are negative score and negative log fold change. Pathway annotations were added by the authors based on data from the Gene Cards database (genecards.org)

| Gene name | neg \| score | neg \| lfc | Pathway annotation |
| --- | --- | --- | --- |
| *FANCL* | 5.71E-15 | -1.9354 | Fanconi Anemia |
| *LMO2* | 9.79E-15 | -1.5989 | Transcription |
| *SLX4IP* | 3.77E-12 | -1.8077 | DNA Repair |
| *BRIP1* | 5.46E-12 | -2.6796 | Fanconi Anemia |
| *UBE2D3* | 9.78E-12 | -1.9414 | DNA Repair |
| *RAD18* | 5.22E-11 | -1.6352 | DNA Repair |
| *ATR* | 8.94E-11 | -2.1514 | DNA Damage Checkpoint |
| *TXNDC17* | 1.39E-10 | -0.88932 | NFKB signaling |
| *RAD1* | 3.00E-10 | -1.0602 | DNA Damage Checkpoint |
| *HELQ* | 1.53E-09 | -1.4685 | Fanconi Anemia |
| *TRIR* | 1.21E-08 | -0.96154 | Telomerase |
| *FANCG* | 2.10E-08 | -1.425 | Fanconi Anemia |
| *FANCA* | 3.53E-08 | -1.587 | Fanconi Anemia |
| *FANCD2* | 6.59E-08 | -1.1835 | Fanconi Anemia |
| *UBE2T* | 7.43E-08 | -1.5685 | Fanconi Anemia |
| *FANCM* | 1.01E-07 | -0.74576 | Fanconi Anemia |
| *DCAF8* | 1.03E-07 | -0.73183 | DNA Damage Response |
| *HUS1* | 1.63E-07 | -0.95024 | DNA Damage Checkpoint |
| *ERCC1* | 3.80E-07 | -0.5416 | Fanconi Anemia |
| *FANCF* | 4.18E-07 | -1.4741 | Fanconi Anemia |
| *WDR26* | 5.41E-07 | -0.74856 | DNA Damage Response |
| *FANCE* | 5.99E-07 | -1.553 | Fanconi Anemia |
| *UBE2H* | 7.76E-07 | -0.83884 | Protein Ubiquitination |
| *BCL2* | 8.30E-07 | -0.82888 | Apoptosis Regulator |
| *SNRNP40* | 9.48E-07 | -0.72294 | Splicing |
| *RAD9A* | 1.13E-06 | -0.98848 | DNA Damage Checkpoint |
| *DYRK1A* | 1.30E-06 | -1.1692 | Homologous Recombination |
| *SLF1* | 1.46E-06 | -0.59398 | DSB repair |
| *UBE2G1* | 1.64E-06 | -0.73631 | Protein Ubiquitination |
| *ERCC5* | 1.65E-06 | -1.0081 | NER, ICL repair |
| *PPM1D* | 1.68E-06 | -1.4666 | Cell Cycle Regulation |
| *FANCB* | 3.18E-06 | -1.0767 | Fanconi Anemia |
| *CPPED1* | 3.28E-06 | -0.95408 | Transcription |
| *EXO1* | 3.40E-06 | -0.50218 | Cell Cycle Regulation |
| *RNF8* | 3.46E-06 | -0.53715 | Cell Cycle Regulation |
| *WSB1* | 4.29E-06 | -0.60328 | Protein Ubiquitination |
| *YPEL5* | 4.31E-06 | -1.0857 | Protein Ubiquitination |
| *FAAP20* | 4.32E-06 | -0.67701 | Fanconi Anemia |
| *TGFBR2* | 4.63E-06 | -0.64834 | Signaling |
| *LYL1* | 5.18E-06 | -0.59682 | Transcription |
| *POLH* | 5.26E-06 | -0.977 | TLS polymerase |
| *RANBP9* | 5.79E-06 | -0.83674 | Protein Ubiquitination, GTPase |
| *XPA* | 6.75E-06 | -0.52743 | NER |
| *ERCC4* | 6.86E-06 | -0.75384 | Fanconi Anemia |
| *SMNDC1* | 7.00E-06 | -0.48524 | Splicing |
| *CCAR1* | 8.04E-06 | -0.70668 | Transcription |
| *hsa-mir-6893* | 8.18E-06 | -0.61157 | Unknown |
| *LIN37* | 8.35E-06 | -0.58991 | Cell Cycle Regulation |
| *RB1CC1* | 9.60E-06 | -0.78433 | DNA Damage Response |
| *FAAP24* | 9.79E-06 | -0.69623 | Fanconi Anemia |
| *INTS15* | 1.03E-05 | -0.72313 | Transcription |
| *FBXO38* | 1.08E-05 | -0.31976 | Protein Ubiquitination |
| *RAD17* | 1.23E-05 | -0.64721 | DNA Damage Checkpoint |
| *GCLC* | 1.24E-05 | -0.69246 | Glutathione Synthesis |
| *NFRKB* | 1.28E-05 | -0.73748 | DNA Damage Checkpoint |
| ***UVSSA*** | **1.47E-05** | **-0.54968** | **TC-NER** |
| *FAAP26* | 1.56E-05 | -0.6257 | Fanconi Anemia |
| *CFAP20* | 1.78E-05 | -0.84912 | Splicing |
| *STRA13* | 1.90E-05 | -0.69879 | Transcription |
| *C1orf112* | 2.33E-05 | -0.62039 | Unknown |
| *FBLN7* | 2.58E-05 | -0.44854 | Cell Adhesion |
| *KIF18A* | 2.59E-05 | -0.47585 | Microtubule |
| *TRAF2* | 2.78E-05 | -0.721 | Apoptosis Regulator |
| *BIRC2* | 3.05E-05 | -0.63634 | Apoptosis Regulator |

**Table S2:** Top SJG-136 induced UVSSA protein-protein interactions. Date from Fig 3D; top 20 proteins significantly enriched in Co-IP of FLAG-UVSSA following SJG-136 treatment, measured by log2 fold change between SJG-136 and mock treated samples. Data is fold change log2(LFQ) and q- value (FDR) for comparison between mock and SJG-136 treated conditions. Annotation performed by authors based on information from the Gene Cards database (genecards.org).

| Gene name | Log<sub>2</sub> FC (LFQ) SJG-136 vs | q-value | Annotation |
| --- | --- | --- | --- |
| *TCOF1* | 3.822508176 | 0.0883401 | DNA damage response |
| *ESCO2* | 3.207551003 | 0.106609579 | Chromatin organization |
| *ZNF706* | 3.130217234 | 0.115014692 | Transcription regulation |
| *ERCC6* | 2.671177228 | 0.106609579 | (CSB) NER |
| *ZFHX4* | 2.114109039 | 0.121836099 | Transcription regulation |
| *COX7A2* | 2.078472137 | 0.106609579 | Mitochondrial protein |
| *ERCC8* | 1.944602648 | 0.132056578 | (CSA) NER |
| *UVSSA* | 1.360626856 | 0.121836099 | (UVSSA) NER |
| *PPRC1* | 0.88415273 | 0.130832447 | Transcription regulation |
| *LTV1* | 0.848051071 | 0.132056578 | Ribosome biogenesis |
| *GTF2H1* | 0.773094177 | 0.130832447 | (TFIIH) NER |
| *GTF2H3* | 0.654779752 | 0.132056578 | (TFIIH) NER |
| *GTF2H4* | 0.637824059 | 0.132056578 | (TFIIH) NER |
| *REXO1* | 0.609376589 | 0.106609579 | Transcription elongation |
| *ERCC3* | 0.597548803 | 0.106609579 | (TFIIH) NER |
| *CENPF* | 0.595184326 | 0.130813074 | Chromatin organization |
| *UFL1* | 0.581530571 | 0.106609579 | DNA damage response |
| *ABCD3* | 0.568094571 | 0.106609579 | Mitochondrial protein |
| *GTF2H2C;GTF2H* | 0.559137026 | 0.121836099 | (TFIIH) NER |
| *VPS72* | 0.538300832 | 0.121836099 | DNA damage response |

## Notes

### Competing Interest Statement

The authors have declared no competing interest.

### Summary of Updates

Results section revised to include new data. Discussion section revised to reference new data and expand discussion of the relevance of the research. Minor text revisions throughout to fit formatting for Molecular Cell and improved readability. Figure 1 revised to streamline panels and update panel 1C with results from a modified experiment. Figure 3 updated to streamline panels and include new data in panel 3F and 3G. Figure S1 revised to support the revisions in figure 1. Figure S2 revised to support revision in figure 3.

